# Spectroscopic DNA-PAINT for simultaneous multiplexed super-resolution microscopy

**DOI:** 10.64898/2026.04.14.718276

**Authors:** Md Abul Shahid, Kripa Patel, David A. Miller, Yang Zhang

## Abstract

Simultaneous multiplexed super-resolution imaging remains a central challenge in single-molecule localization microscopy (SMLM), due to the limited number of spectrally distinguishable fluorophores and the trade-off between spatial and spectral precision. Here, we introduce spectroscopic DNA-PAINT (sDNA-PAINT), a framework that integrates DNA-PAINT with spectroscopic SMLM (sSMLM) to enable simultaneous multiplexed imaging with high spatial and spectral fidelity. Using DNA Origami Nanorulers, *in vitro* and fixed-cell imaging, we show that sDNA-PAINT conditions significantly improve spectral precision and photon budgets compared to conventional glass and antibody-conjugated conditions in sSMLM imaging. Across representative dyes from three fluorophore families (Rhodamine, Cyanine and Oxazine), narrow spectral centroid distributions are observed to enable reliable statistical discrimination at the single-molecule level, even for dyes with heavily overlapping ensemble spectra. In dual-target cellular imaging, sDNA-PAINT achieves accurate spatial reconstruction and high classification accuracy, demonstrating high-fidelity simultaneous multiplexing within a single acquisition. sDNA-PAINT provides a pathway toward high-throughput simultaneous multiplexed super-resolution interaction imaging.

---

Single-molecule localization microscopy (SMLM)^1,2^ technologies such as photoactivated localization microscopy (PALM)^3–7^, stochastic optical reconstruction microscopy (STORM)^8–11^, and Points Accumulation for Imaging in Nanoscale Topography (PAINT)^12,13^, have transformed the visualization and spatially-resolved investigation of nanoscale biological systems by achieving sub-diffraction resolution, single-molecule sensitivity, and multicolor imaging. These techniques rely on fluorescence intermittency, in which individual fluorophores stochastically switch between “on” and “off” states, enabling precise localization of single molecules at the nanoscale.^1^ Recently developed spectrally resolved STORM (SR-STORM)^14–21^or spectroscopic SMLM (sSMLM, **Figure 1a**)^22–28^, pioneered by others and by our group, enables the concurrent acquisition of both spatial and spectral information of individual fluorescent molecules. By statistically discriminating single-molecule emission spectra from different fluorophore species, sSMLM allows simultaneous multiplexed imaging even in the presence of substantial spectral overlap. However, a major challenge lies in identifying a set of fluorophores that are both spectrally distinguishable and compatible under a single set of imaging conditions. To date, the only well-established fluorophore sets that meet these criteria are conventional STORM-based far-red cyanine dyes.^19,26^ Despite their widespread use, their highly variable photobleaching kinetics and photon budgets limit performance and consistency (right panel in **Figure 1b**).^9^ In addition, because photons must be divided between spatial and spectral detection channels, sSMLM inherently suffers from reduced spatial resolution.^29^ This limitation is further compounded by <u>s</u>ingle-<u>m</u>olecule <u>flu</u>orescence <u>s</u>pectral <u>h</u>eterogeneity (smFLUSH)^30^, which refers to variations in emission spectra among individual molecules of the same fluorophore across different locations within a sample. Together with limited photon counts in the spectral channel, these factors reduce spectral precision^29^, constrain molecular identification accuracy, and ultimately limit the number of resolvable color channels.

**Figure 1.**
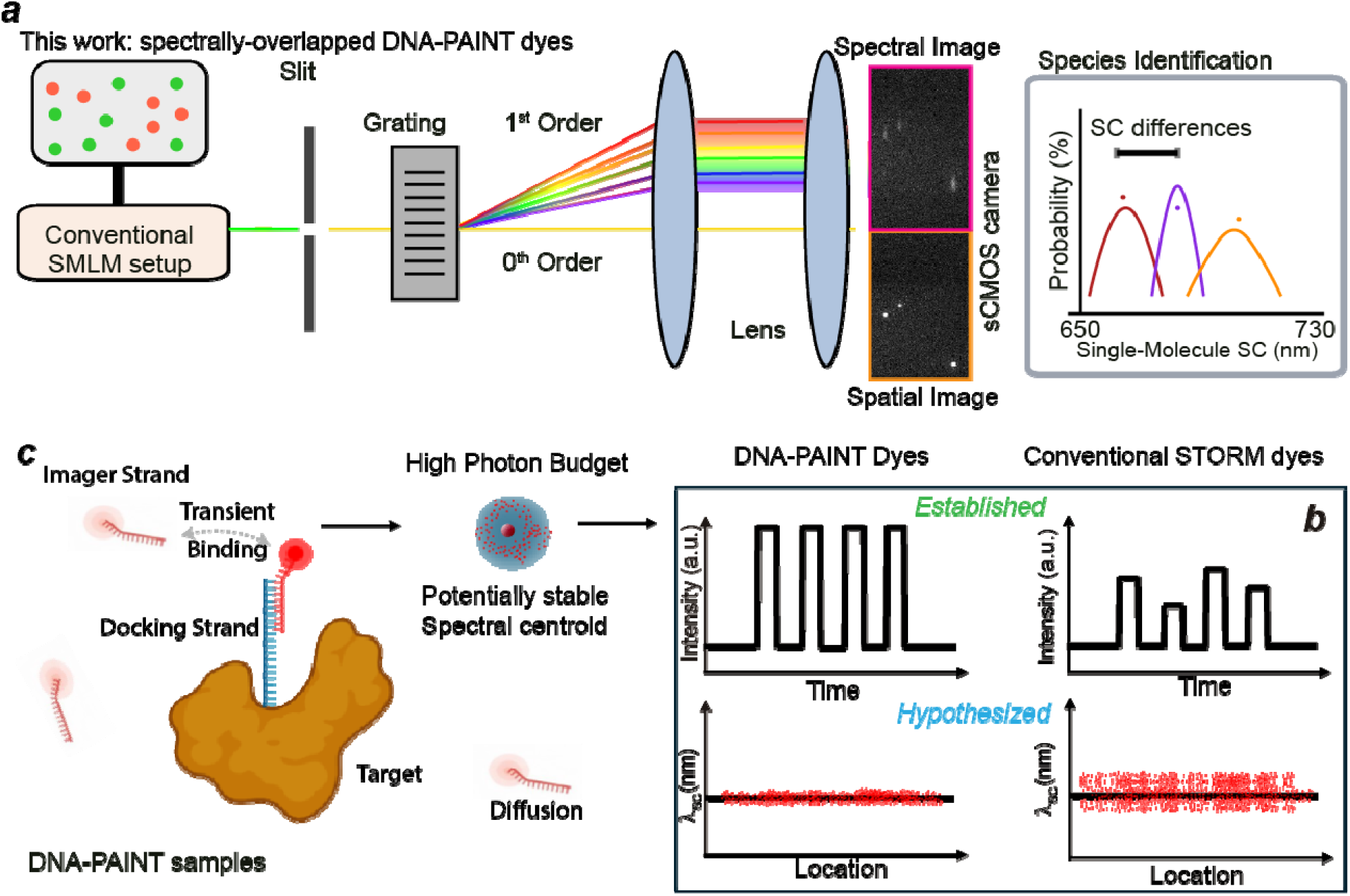
(***a***) schematic of an sSMLM system and the single-molecule discrimination based on probability distribution of SCs; (***b***) the hypothesized benefits of DNA-PAINT in achieving high-photon budget and high single-molecule spectral homogeneity in comparison to conventional STORM-based dyes; **(c)** Illustration of DNA-PAINT principle

Over the past decade, DNA-based PAINT (or DNA-PAINT) has emerged for advancing SMLM toward true molecular resolution (< 2 nm).^31–40^ This capability stems from the programmable and transient hybridization between short “docking” strands, covalently attached to biomolecular targets (*e.g*., antibodies), and complementary, fluorophore-labeled “imager” strands in solution (**Figure 1c**).^35^ The imager strands freely diffuse until transient binding occurs, generating fluorescence bursts lasting tens to hundreds of milliseconds and yielding up to 10□ detected photons per event^41^, which enables exceptionally high localization precision. Moreover, the orthogonality of DNA sequences allows multiple targets to be encoded with distinct docking strands, enabling highly multiplexed imaging through sequential introduction of corresponding imager strands, as demonstrated in Exchange-PAINT.^33,42,43^

A method that enables the parallel acquisition of multiple DNA-PAINT targets under sSMLM imaging setup would address the limitation for sSMLM to improve the spatial resolution in comparison with conventional sSMLM/SR-STORM. At the same time, the integration could reduce acquisition time for any sequential SMLM/DNA-PAINT proportionally to the number of simultaneously imaged channels. Therefore, we introduce spectroscopic DNA-PAINT (sDNA-PAINT), a new framework for simultaneous multiplexed super-resolution imaging. The key hypothesis underlying our approach is that the single-molecule photophysical behavior of fluorophores under DNA-PAINT conditions is particularly well suited for single-molecule spectral discrimination. First, as DNA-PAINT decouples single-molecule blinking from fluorophore photochemistry and photophysics by transient hybridization, it directly opens up th choice of more fluorophore families for simultaneous multiplexed sSMLM to achieve high and programmable photon budget (**Figure 1b** left panel). Second, we hypothesize that smFLUSH, often arising from local environmental variability, is suppressed in DNA-PAINT. This feature, together with the inherently high photon budget (and reduced noisy-uncertainty) in DNA-PAINT system collectively result in lower spectral uncertainty^29^ for sSMLM system (**Figure 1b** left panel). Here spectral uncertainty is defined as the standard deviation (σ_SC_) of the observed single-molecule intensity-weighted spectral centroid (*λ*_SC_). Presumbably, imager strands freely diffuse in solution and remain conformationally flexible upon transient binding to their targets. This dynamic and relatively homogeneous solvation process presumably reduces local perturbations to the fluorophore, promoting a more uniform emission spectrum across individual molecules. To investigate these benefits in integrating DNA-PAINT and sSMLM, we systematically characterized the photon budget and smFLUSH properties of four commonly used DNA-PAINT fluorophores and compared their spectral behavior to that of conventional antibody-dye conjugates and on glass substrates. Our results show that DNA-PAINT imaging conditions yield relatively stable and homogeneous spectral profiles in addition to the well-known brighter and programmable photon counts, providing a critical foundation for robust and accurate spectral unmixing and localization at sub-10 nm level.

To evaluate the performance of sDNA-PAINT and quantify single-molecule spectral behavior, we first investigated the suitability of implementing DNA-PAINT method on a sSMLM optical platform. The detailed optical setup of our sDNA-PAINT/sSMLM system was indicated in **Figure S1**. To begin, we imaged a DNA origami nanoruler sample obtained from GATTAquant as an reference standard.^44^ The illumination power and exposure time were systematically investigated to provide an optimal localization precision, photon budget at an exposure time of 150 milliseconds (ms) and a power density of 1.4 kW cm^-2^ (**Figure S2**). The full field-of-view (FOV) shows well-dispersed nanorulers (**Figure 2a**), and magnified views visualize the three docking sites of two distinct nanorulers within the individual structures (**Figure 2b**). All samples were acquired under the total internal refraction fluorescence (TIRF) angle unless otherwise noted. The measured localization pattern matches the designed geometry, with three binding sites separated by ∼80 nm (**Figure 2c)**. Specifically, line profile analysis across an individual DNA nanoruler confirms this spacing, giving separations of 80.9 nm and 78.6 nm among the three dots (**Figure *2b-c***), which indicates high spatial resolution under these imaging conditions with an average localization uncertainty of ∼7 nm (**Figure S3**).

**Figure 2.**
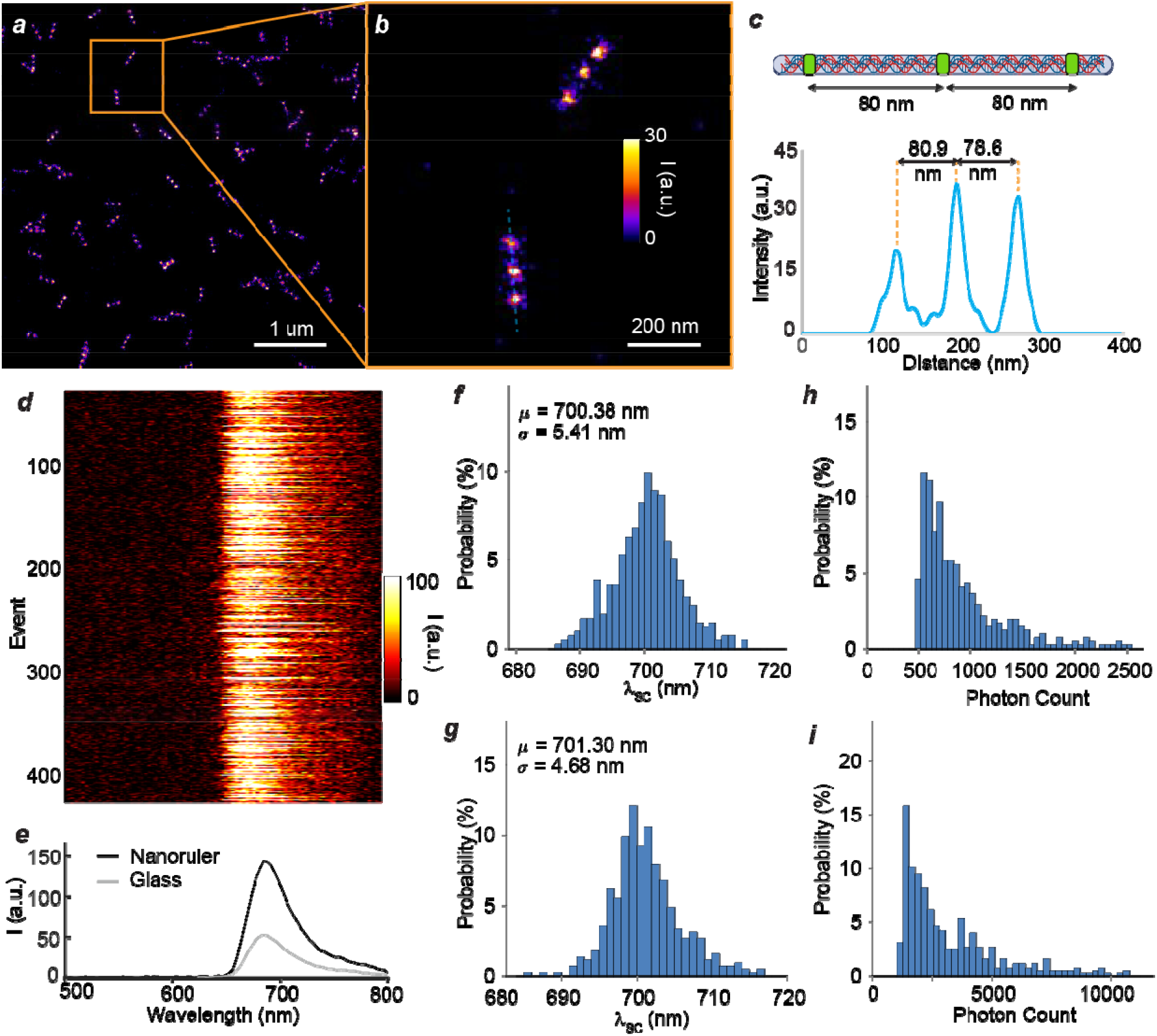
(***a-b***) **s**DNA-PAINT image of DNA Origami Nanoruler: Full FOV (***a***) and the magnified views (***b***) of the rectangular box in ***a***, (***c***) top: the illustration of 80-nm spaced molecular labeling position on Nanoruler and bottom: the line profile of the experimentally measured results of a nanoruler from the dashed line in ***b***; (***d***) representative 454 single-molecule emission spectra of ATTO655 measured on from Nanoruler sample; (***e***) averaged single-molecule emission spectrum from Nanoruler (black curve) and bare glass substrate (gray curve); (***f-i***) Histograms of *λ*_SC_ (***f-g***) and photon counts (***h-i***) for ATTO655 measured on the glass substrates (***f*** and ***h***) and under Nanoruler (***g*** and ***i***).

We then examined the spectral properties of single molecules under sDNA-PAINT conditions. A total of 454 individual emission spectra of ATTO655 were collected from nanoruler-bound imager strands (**Figure 2d**). High-throughput single-molecule spectroscopy (HT-SMS) was performed by reducing the entrance slit width to 100 um, adapting our previously established setup^30^ to minimize noise contributions to spectral uncertainty. The averaged single-molecule spectrum closely matches the reference spectrum measured on a bare glass substrate and reported solution-measurement value^45^, indicating that the sDNA-PAINT environment does not introduce noticeable spectral change (**Figure 2e**). We further analyzed the probability density distributions of the λ_SC_ and photon counts. The single-molecule λ_SC_ distributions measured on glass and on nanorulers (**Figure 2f–g**) show means of 700 nm and 701 nm as well as and σ_SC_ 5.41 and 4.68 nm, respectively. The photon count (**Figure 2h–i**) measured on bare glass substrate (mean = 922) is significantly lower than that measured on DNA-PAINT condition (mean = 2620) which is expected because of the programmable and high photon budget in the latter case. We note that the photon counts are calculated only from the spectral channel with about 50% overall transmission efficiency to the first-order diffraction in our sSMLM system. These results indicate that the sDNA-PAINT system maintains accurate spatial reconstruction while achieving lower □_SC_, supporting reliable spectral discrimination for multiplexed imaging.

Next, we investigated the smFLUSH properties in fixed BSC-1 cells labeled separately with different dyes used in DNA-PAINT conditions. We intentionally selected four dyes from three fluorophore families (*i.e*., Rhodamine/ATTO565, Cyanine/Cy3B, Oxazine/ATTO655 and ATTO680) to investigate the generatability of our approach. The dyes are synthetically attached on the respective imager strands and obtained from Massive Photonics. The cells were immunostained with antibodies attached to the complementary docking strand and sDNA-PAINT imaging was performed in the same imaging buffer and acquisition conditions as **Figure 2** (see detailed imaging protocol in **Supporting Information**). Single-color sDNA-PAINT reconstructions of mitochondria and vimentin fibers showed well-defined subcellular structures consistent with expected worm-like elongated (**Figures 3a and 3c**) and fiber-like morphologies (**Figures 3b and d**). Spectroscopically, all dyes exhibited stable emission spectra with narrowly distributed λ_SC_ (**Figures 3e-h**) with □_SC_ of 2.98, 1.95, 3.15 and 3.21 nm respectively for ATTO565, Cy3B, ATTO655 and ATTO680 (**Table 1**), indicating relatively low spectral heterogeneity across localizations throughout the entire FOV.

**Table 1.** Ensembled and single-molecule photophysical properties of tested dyes under three imaging conditions.^a^.

| | $\lambda_{Em}^b$ | $\lambda_{SC}$ (DNA-PAINT)<sup>c</sup> | $\lambda_{SC}$ (Glass)<sup>c</sup> | $\lambda_{SC}$ (Antibody)<sup>c</sup> |
| --- | --- | --- | --- | --- |
| ATTO565 | 593 | 610.1±2.98 | 612.6±5.32 | 607.9±9.69 |
| Cy3B | 580 | 603.2±1.95 | 611.5±5.37 | 609.7±10.64 |
| ATTO655 | 682 | 697.8±3.15 | 689.2±4.89 | 693.4±5.18 |
| ATTO680 | 702 | 713.5±3.21 | 707.38±5.89 | 708.3±3.86 |

<sup>a</sup>all unit reported in nm; <sup>b</sup>averaged single-molecule emission spectral peak measured using sDNA-PAINT; <sup>c</sup> the ± number indicates the mean and standard deviations of the $\lambda_{SC}$ measured among >500 individual molecules in each condition

**Figure 3.**
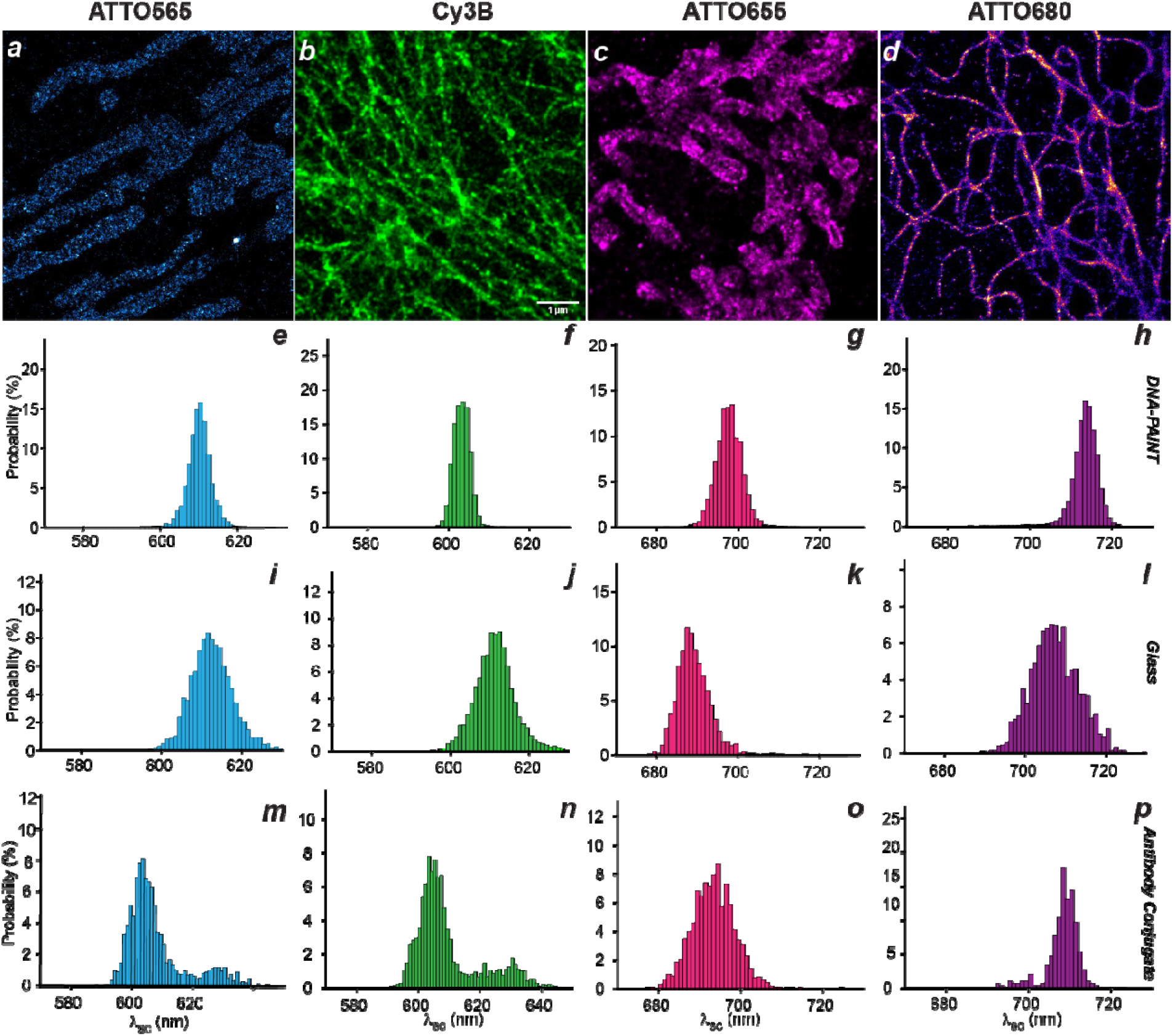
sDNA-PAINT imaging and smFLUSH results on different conditions. (***a-d***) Single-color sDNA-PAINT reconstructed images using ATTO565, Cy3B, ATTO655 and ATTO680 labeled on mitochondria (***a, c***) and vimentin fibers (***b, d***) in fixed BSC-1 cells; (***e-p***) the corresponding histograms of probability distributions of λ_SC_ measured under DNA-PAINT conditions (***e-h***), bare glass substrates (***i-l***) and covalently attached to antibodies (***m-p***) for ATTO565, Cy3B, ATTO655 and ATTO680, respectively.

For side-by-side comparison, we measured the same dyes deposited on bare glass substrates. In this condition, all dyes showed broader λ_SC_ distributions (**Figure 3i–l**) with increased □_SC_ ranging from 4.68nm – 5.89 nm. The increased intrinsic smFLUSH of the deys under nanoscale heterogeneous environment on the glass surface likely has a major contribution to the observed larger σ_SC_ since the pure noise-uncertainty contribution in our sSMLM system under this condition is < 2 nm according to numerical simulations.^30^ We further evaluated the dyes when covalently conjugated to secondary IgG antibodies, the most common labeling strategy in SMLM^8,19^. The antibody conjugates were prepared with a degree of label <1.3 per antibody to prevent dye aggregation (**Table S1**). Under these conditions, all four dyes exhibited substantially increased □_SC_ up to 10.64 nm (**Figure 3m–p**), despite maintaining similar mean λ_SC_ values compared to DNA-PAINT (**Table 1**). The only exception is for ATTO680 which possesses lower σ of 3.86 nm versus 5.89 nm on glass. The increase in smFLUSH suggests that the local environment associated with protein conjugation introduces additional variability in single-molecule emission behavior. This variability presumably arises from site-non-specific labeling at different amino acid residues from the N-hydroxysuccinimide (NHS)-coupling reaction^46^, which can modulate the local conformation and alter the excited-state dynamics of individual fluorophores. In short, these results indicate that DNA-PAINT provides a more uniform photophysical and local environment that minimizes smFLUSH and spectral uncertainty. This property is critical for sSMLM, where accurate spectral discrimination depends on minimizing overlap between single-molecule spectral distributions.

Finally, with the smFLUSH characteristics and statistical separability of individual dyes established, we next performed simultaneous multiplexed sDNA-PAINT imaging in fixed cells. From the separately acquired histograms, four dyes can be spectrally resolved with statistical confidence (**Figure 4a**). For example, the ATTO655 and ATTO680 exhibited distinct single-molecule emission profiles, with average emission peaks at 682 nm and 702 nm and λ_SC_ means of 697.8 nm and 713.5 nm, respectively. The λ_SC_ distributions remained narrow, with σ_SC_ of 3.15 nm and 3.21 nm, enabling reliable assignment of individual localizations to their corresponding targets. Similarly, Cy3B and ATTO565 could be statistically resolved based on the individual smFLUSH characterization. To evaluate the theoretical discriminability of the two fluorophores based solely on λ_SC_, we approximated their single-molecule λ_SC_ distributions as Gaussian functions using the experimentally measured means and standard deviations. Under this model, the separation between the two distributions (∼15.7 nm for ATTO655/ATTO680) is approximately five times larger than their σ_SC_ (∼3 nm). Estimating the probability that molecules from one distribution fall within the ±1σ range of the other yields cross-misidentification rates on the order of 10^-5^ (**Note S1** in **Supporting Information**). These results indicate negligible overlap and confirm that SC alone is sufficient for robust discrimination for this dye pair.

**Figure 4.**
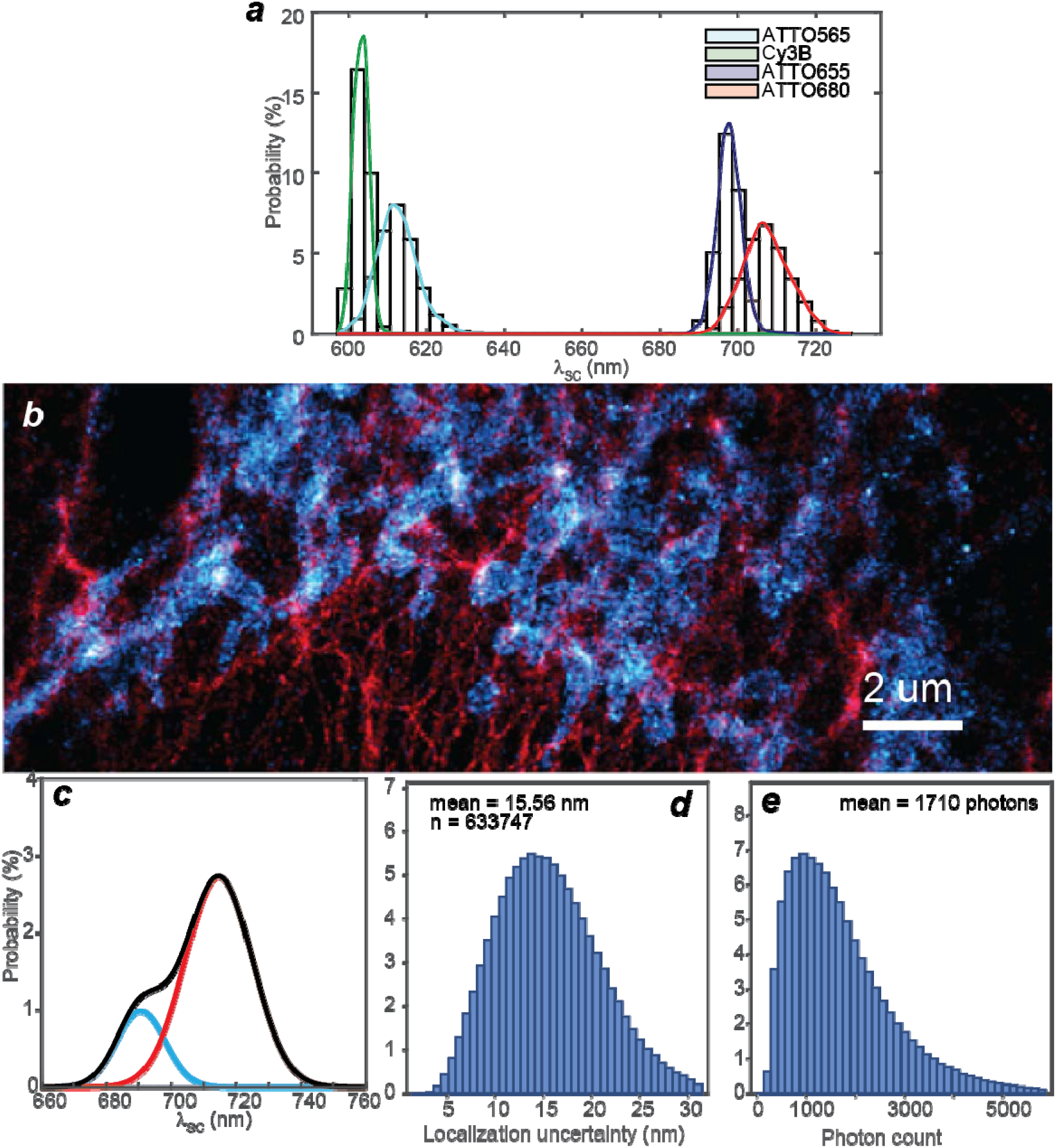
(***a***) Histograms of probability distributions of λ_SC_ of ATTO565, Cy3B, ATTO655, ATTO680 measured separately under DNA-PAINT conditions in fixed BSC-1 cells; (***b***) Two-color simultaneou sDNA-PAINT image of mitochondria (blue) and vimentin fiber (red) of a fixed BSC-1 cell labeled with ATTO655 and ATTO680, respectively; (***c-e***) Histograms of probability distribution of λ_SC_ (***c***), localization precision (***d***), and photon counts (***e***) among 633,747 single molecules detected in ***b*** together with a two-component gaussian fits to the histogram in ***c***.

Following the above statistical confidence that the two DNA-PAINT dyes can be statistically resolved in sSMLM with <0.01% misidentification rate^27^, we then acquired simultaneous two-color sDNA-PAINT imaging using ATTO655 and ATTO680 to label mitochondria and vimentin within the same cell. Only single molecules with λ_SC_ falling into the range of 694.65 nm –700.95 nm and 710.29 nm and 716.71 nm were collected and classified as ATTO655 and ATTO680 single-molecule localizations color-coded in blue and red respectively (**Figure 4b**). The rendered two-color sDNA-PAINT image reveals clear worm-like structure in the blue (ATTO655) channel and fibers in the red (ATTO680) channel, clearly demonstrating accurate spectral classification of the two distinct structures. The histogram of λ_SC_ values across all detected molecules shows two well-separated populations that are well described by a two-component Gaussian fit (**Figure 4c**) peaked at 690.6 nm and 714.7nm, which agree well with the separately measured λ_SC_ of 697.8 nm and 713.5 nm for ATTO655 and ATTO680, respectively. The average localization precision obtained from this image is 15.6 nm with a photon budget around 1710 (**Figure 4d-e**). The localization precision is improved by 40% compared with conventional simultaneous multiplexed sSMLM labeled with cyanine dyes using the identical optical setup.^17^

Above results show that the reduced smFLUSH observed in DNA nanorulers and single-target experiments is preserved in multiplexed sDNA-PAINT imaging. The DNA-mediated transient binding environment produces high photon output and stable spectral signatures, which reduces spectral uncertainty and reduces overlap in the histograms between dye populations. In contrast, protein-conjugated dyes exhibit broader spectral distributions and lower classification accuracy under comparable conditions. This behavior addresses a central challenge in multiplexed sSMLM, where reliable classification depends on both photon budget and spectral stability. DNA-PAINT conditions increase photon yield, reduce smFLUSH, and maintain consistent spectral peak positions across large numbers of binding events. These factors improve effective spectral resolution, even for dyes with modest ensemble spectral separation. As a result, DNA-PAINT supports high-fidelity simultaneous spectral multiplexing of sSMLM within a single acquisition.

Despite these advantages, several limitations remain on both the instrumentation and probe sides. In our current sSMLM imaging setup, the grating we used was specifically selected for far-red channel imaging with 1:3 splitting ratio between the spatial and spectral channel to provide sufficient photons for spatial localization and spectral analysis. However, in the green channel the splitting ratio is about 1:20^47^, which prevents good localizations for Cy3B and ATTO565 under DNA-PAINT condition. Other reported methods based on prism/beam splitters^19^ and dual-wedge prisms^48^ could potentially push the system towards >6-color simultaneous multiplexing at the expense of more complicated optical setup and non-linear spectral dispersion. Alternative detection schemes in sSMLM/SR-STORM could further enhance performance when combined with the stabilized photophysical environment provided by DNA-PAINT. For example, biplane SR-STORM^13^ and symmetrically dispersed spectral detection architectures^19^ can increase photon utilization efficiency by doubling photon collection or conserving photons between spatial and spectral channels. These approaches can improve localization precision and spectral accuracy without sacrificing multiplexing capability while being fully compatible with our proposed sDNA-PAINT framework. The dyes tested in this study are primarily limited to established DNA-PAINT fluorophores but can be readily extended to other advanced, bright, and photostable fluorophores, including boron dipyrromethenes (BODIPYs) and self-healing dyes.^6,49^

In conclusion, this work establishes sDNA-PAINT as an effective framework for simultaneous multiplexed super-resolution imaging by integrating the programmable binding of DNA-PAINT with sSMLM. Systematic measurements using DNA origami nanorulers, *in vitro* and fixed-cell imaging, and dual-target experiments show that DNA-PAINT conditions produce stable and homogeneous single-molecule spectral signatures while maintaining high spatial accuracy. In general, under the sDNA-PAINT condition, the smFLUSH (2-3 nm) reduced by ∼50% from 4-5 nm in glass and 4-10 nm in antibody-conjugated conditions. The reduced smFLUSH and increased photon output enable improved spectral precision, which directly supports reliable classification of individual molecules even when fluorophores exhibit overlapping ensemble spectra. Comparisons across DNA-PAINT, bare glass, and antibody-conjugated conditions reveal that the local molecular environment plays a central role in determining spectral variability. DNA-PAINT provides a controlled and reproducible binding context that minimizes spectral uncertainty and preserves consistent emission characteristics across different dyes and cellular targets. This behavior enables robust multiplexed imaging without relying on large spectral separation between fluorophores. The results point to a broader design strategy for spectrally-resolved super-resolution imaging. Instead of focusing solely on fluorophore development, engineering the single-molecule environment can significantly improve spectral discriminability. DNA-PAINT offers a practical implementation of this concept, supporting higher effective spectral resolution and expanding the number of distinguishable targets within a single acquisition. sDNA-PAINT provides a path toward high-fidelity, multi-target imaging with reduced acquisition time and improved reconstruction fidelity. The ability to achieve accurate spatial localization and reliable spectral identification within the same experiment creates new opportunities for studying complex biological systems and interactions at the nanoscale.

## ASSOCIATED CONTENT

### Supporting Information

Imaging setup and procedures; smFLUSH characterization; antibody conjugations; statistical analyses. Following files are available free of charge. brief description (file type, i.e., PDF)

## AUTHOR INFORMATION

### Notes

D.A.M. serves as a consultant for Lumedica Vision Inc. This affiliation had no role in the funding or execution of this work. The other authors declare no competing financial interests.

## ACKNOWLEDGMENT

Y.Z. acknowledges support from the National Science Foundation (CHE-2246548 and CHE-2441081), the National Institutes of Health (R21GM141675 and R35GM155241). M.A.S and K.P. acknowledge the support of the North Carolina State University Comparative Medicine Institute.

## Notes

### Competing Interest Statement

The authors have declared no competing interest.

