## Supporting Information for "Spectroscopic DNA-PAINT for simultaneous multiplexed super-resolution microscopy"

### 1. sSMLM/sDNA-PAINT optical setup

The emission signals of the single-molecule emitters were collected by the TIRF objective, passed through a dichroic mirror and a long-pass filter, and focused by a tube lens to an intermediate imaging plane, where an entrance slit was set. A relay optics system with a 100 mm focus lens was used to extend the image further, and an EMCCD camera (Teledyne Photometrics 95B or Andor iXon 897) was placed at the focal plane of the second relay lens. A transmission grating (Star Analyser 100, Groove density = 100 lines/mm) was placed between the entrance slit and the first relay lens, and its position along the optical axis could be adjusted to achieve different spectral dispersions (or spectral resolutions) ranging from 2 to 7 nm/pixel. The width of the entrance slit was adjusted to minimum field of view of spatial channel images, enabling high-throughput single-molecule spectroscopy (HT-SMS) with massive scanning and preventing the overlapping of single molecules in spatial channel.^3^

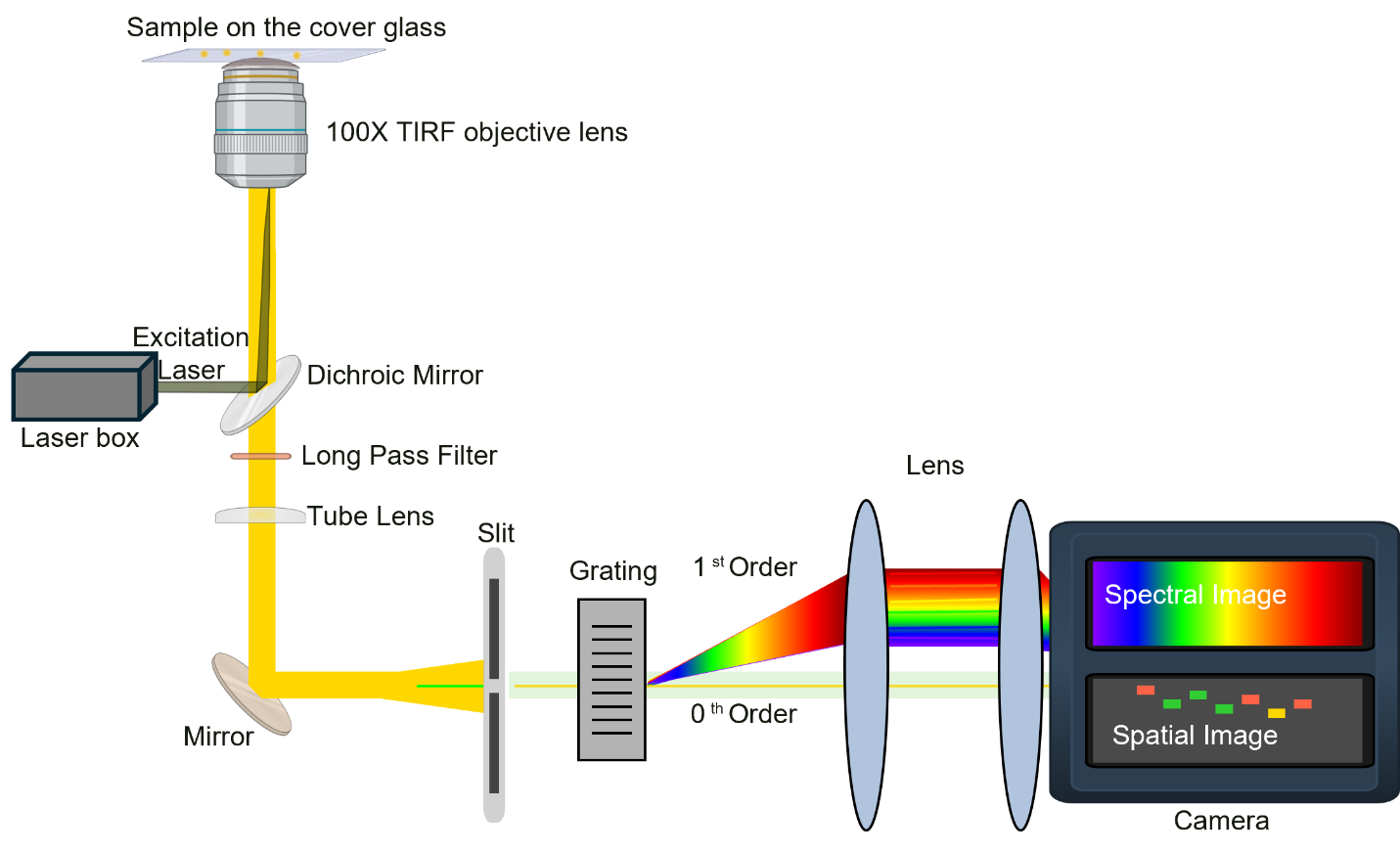

**Figure S1** A schematic of our custom-built sSMLM imaging system, which is constructed from a Nikon Ti-2E inverted fluorescence microscope equipped with a total internal reflection fluorescence (TIRF) arm that allows for switching between wide-field and TIRF configurations. High-power continuous-wave lasers with wavelengths and powers of 488, 561 nm and 640 nm (200 mW, 200 mW, and 500 mW respectively) (Oxxius) were combined and coupled to the single-mode fiber and sent to the input port of the TIRF arm of the Nikon microscope. The laser beam is focused on the back focal plane of a 100 × TIRF objective (Nikon CFI SR HP APO 1.49 OIL). During experimental acquisition, the illumination angle was adapted to TIRF/critical angle to illuminate the dyes at the bottom of the cover glass chamber

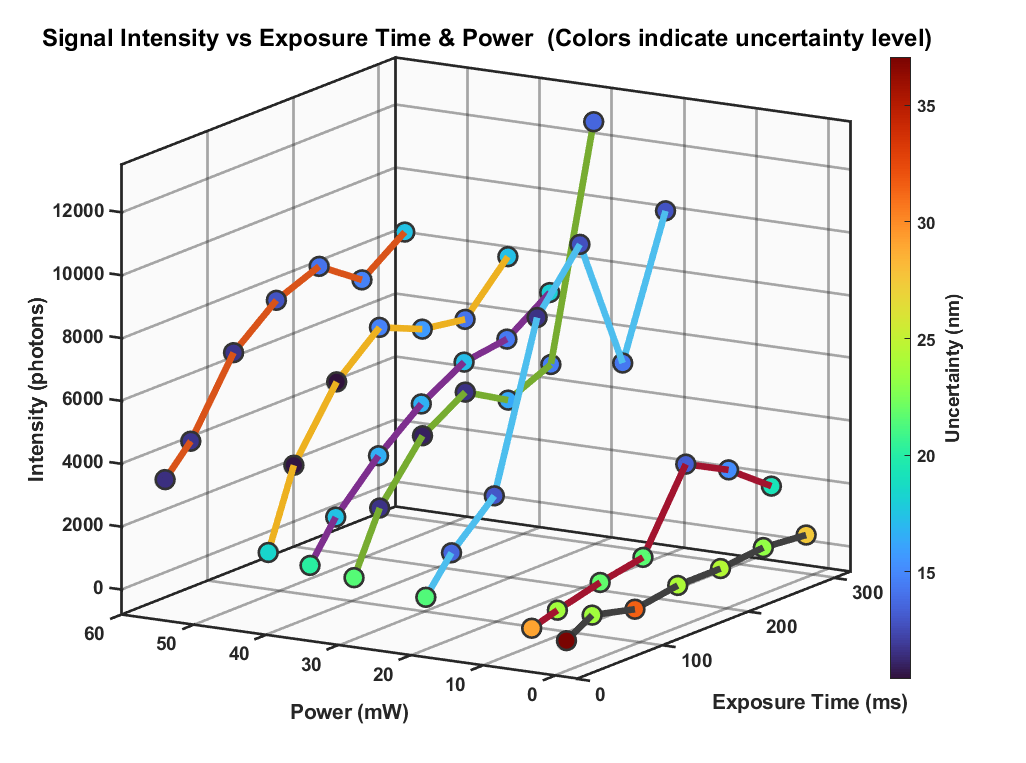

**Figure S2.** Relationship between power, localization precision and exposure time.

To identify optimal acquisition parameters for DNA-PAINT imaging, we systematically varied the excitation laser power and camera exposure time using ATTO655-labeled nanoruler samples (GATTA-PAINT 80R, GATTAquant). Seven discrete laser power levels were tested using an adjustable neutral-density filter wheel (Thorlabs, #88-398, position #1 through #10, corresponding to powers ranging from 2.2 kW cm^-2^ – 28.9 W cm^-2^), each at seven exposure times (20, 50, 100, 150, 200, 250, and 300 ms), yielding a total of 49 acquisition conditions. For each condition, the mean single-molecule signal intensity (photons) and uncertainty (nm) were quantified from localized binding events (Figure S2). Signal intensity scaled approximately linearly with both laser power and exposure time, as expected. At low power, exposure combinations, insufficient photon counts led to elevated uncertainty (>30 nm), while excessively high-power exposure combinations approached detector saturation without further improvement in precision. Balancing high photon yield against minimal uncertainty, we selected filter #4 (~30 mW) with a 150 ms exposure time as the optimal operating condition for all subsequent DNA-PAINT experiments. This regime provided robust single-molecule intensity (~5,000 - 7,000 photons) with uncertainty below ~15 nm, ensuring reliable spectral centroid determination imaging. With further filtering off the spots out of focus and dimmer spots, we achieved the localization uncertainty histogram in **Figure S3**.

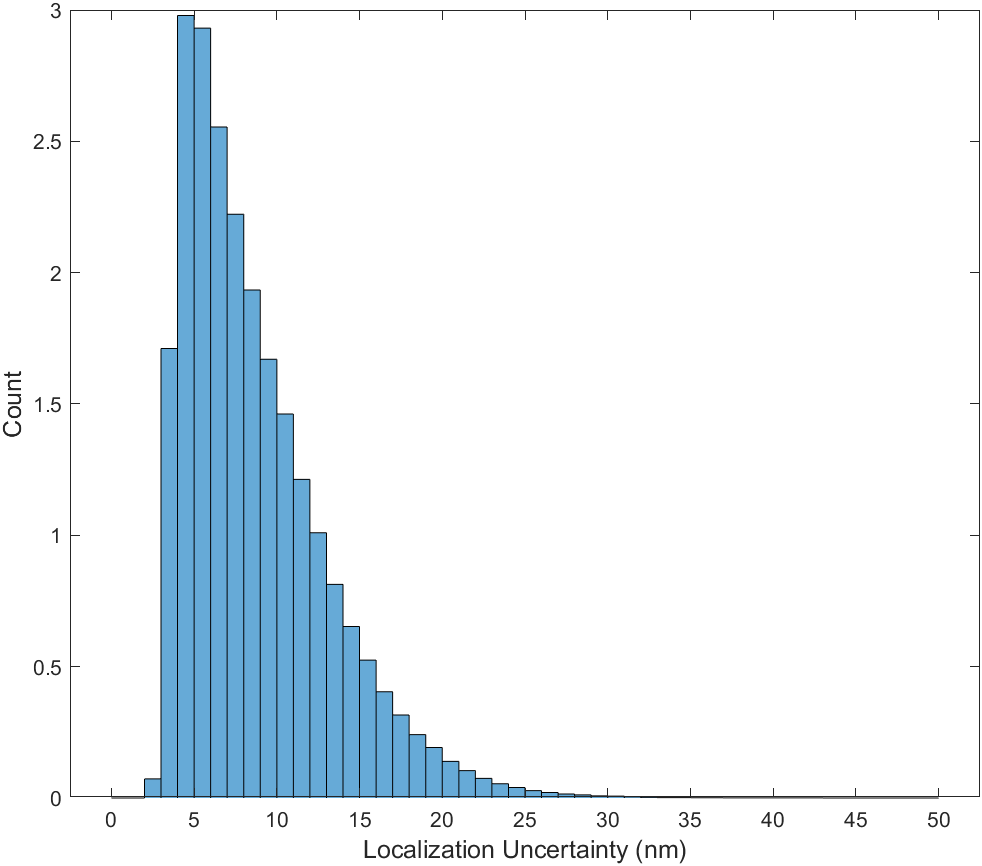

**Figure S3**. Localization Uncertainty of nanoruler DNA-PAINT imaging in Figure 2a

**Note S1. Calculation of statistical confidence based on individually measured single-molecule spectral centroid distributions**

To quantify the discriminability of two fluorophores using only the single-molecule spectral centroid (λ_SC_), we modeled their λ_SC_ distributions as Gaussian random variables based on experimental measurements. The two dyes are characterized by mean SC values of μ₁ = 697.8 nm and μ₂ = 713.5 nm, with corresponding standard deviations σ₁ = 3.15 nm and σ₂ = 3.21 nm.

**Gaussian approximation**

We assume:

$$X_{1}\mathcal{\sim N(}\mu_{1},\sigma_{1}^{2}),X_{2}\mathcal{\sim N(}\mu_{2},\sigma_{2}^{2})$$

The separation between the two means is:

$$\Delta\mu=\mu_{2}-\mu_{1}=15.7\text{ nm}$$

which is approximately five times larger than the standard deviations (~3 nm), indicating minimal expected overlap.

**Misidentification under ±1σ filtering**

To estimate practical misclassification, we consider only molecules within one standard deviation of each mean:

- Dye 1 acceptance window:

$$[\mu_{1}-\sigma_{1},\text{ }\mu_{1}+\sigma_{1}]=[694.65,\text{ }700.95]\text{ nm}$$

- Dye 2 acceptance window:

$$[\mu_{2}-\sigma_{2},\text{ }\mu_{2}+\sigma_{2}]=[710.29,\text{ }716.71]\text{ nm}$$

Misidentification occurs when a molecule from one distribution falls within the acceptance window of the other.

**Probability estimates**

For dye 1 being misidentified as dye 2:

$$P(710.29\leq X_{1}\leq716.71)\approx3.5\times{10}^{-5}$$

For dye 2 being misidentified as dye 1:

$$P(694.65\leq X_{2}\leq700.95)\approx4.0\times{10}^{-5}$$

These probabilities are obtained by evaluating the tails of the corresponding Gaussian distributions.

**Implications**

The extremely low overlap between the two distributions demonstrates that, for this dye pair, spectral centroid alone provides highly reliable discrimination. This analysis highlights that when centroid separation substantially exceeds spectral heterogeneity, simple statistical separation is sufficient without requiring additional spectral features or advanced classification methods.

### 2. Sample preparation

#### 2.1. Dye-antibody conjugation

A secondary antibody (100 uL, 1.3 mg/mL, AffiniPure® Donkey Anti-Rabit IgG (H+L), 715-005-152, JacksonImmuno) was mixed with ~1 uL of the NHS-ester versions of Cy3B (Fisher Scientific), ATTO565, ATTO655, ATTO680 (Sigma) and 10 uL of 1 M NaHCO_3_ for 12 h at room temperature under dark and gentle rocking. The dye-antibody conjugates were then purified using 0.5 mL Amicon Ultra Centrifugal Filters with 50 Kda MWCO and diluted to a protein concentration of 50 ug/mL as stock solutions and refrigerated and used within two weeks. The concentration of the proteins are measured using Nanodrop One spectrometer (Fisher).

#### 2.2. Dye samples deposited on bare glass substrates

Methanol solutions of dyes **(**100 nM, 200 μL) were spin-coated onto bare glass substrates (#1 premium cover glass, 24 × 40 mm^2^, Fisher), using a EZ4 programmable Spin Coater for 1 min at 2000 rpm, dried under vacuum and mounted onto the microscope stage to be imaged on the same day.

#### 2.3. Dye-antibody conjugate samples deposited on bare glass substrates

Dye–antibody conjugates were diluted in imaging buffer to a final concentration of 0.01 µM. A volume of 20 µL of the diluted conjugate was added to each well of an 8-well chamber (Lab-Tek II chambered #1.5 German Coverglass system) and incubated for 15 minutes to allow adsorption of the molecules onto the glass surface. Additionally, a clean glass was placed top of the solution to make a sandwiched system. Following incubation, imaging was performed under TIRF conditions to selectively excite fluorophores at the surface.

**Table S1.** Measurement and characterization of Degree of Labeling of the prepared dye-antibody conjugates

| Dye name | A280 | A (Dye) | ε (M^-1^) | IgG (mg/mL) | Dye (µM) | IgG (µM) | DOL |
| --- | --- | --- | --- | --- | --- | --- | --- |
| ATTO 655 | 1.215 | 0.587 (656 nm) | 125000 | 0.890 | 4.7 | 5.6 | 0.84 |
| Cy3B | 2.026 | 1.526 (559 nm) | 130000 | 1.451 | 11.7 | 9.1 | 1.29 |
| ATTO 565 | 1.960 | 0.393 (568 nm) | 120000 | 1.457 | 3.3 | 9.1 | 0.36 |
| ATTO 680 | 2.036 | 0.636 (681 nm) | 125000 | 1.469 | 5.1 | 9.2 | 0.55 |

### 3. Imaging Acquisition and Analysis Procedure for smFLUSH

The spectroscopic response of this imaging spectrometer was calibrated with a fluorescence lamp before each imaging session; the spectral dispersion is 3.5nm/pixel in this experiment. The single-molecule images were acquired with a camera exposure time of 150 ms, and around 1000 single molecules were acquired in each run. After the acquisition, the single-molecule images were processed with customized codes in MATLAB 2023b. All pure dyes and dye-antibody conjugates samples were excited at 561 or 640 nm with a power density of 1.2 kW/cm² and at 561 nm with a power density of 1.1 W/cm², respectively.

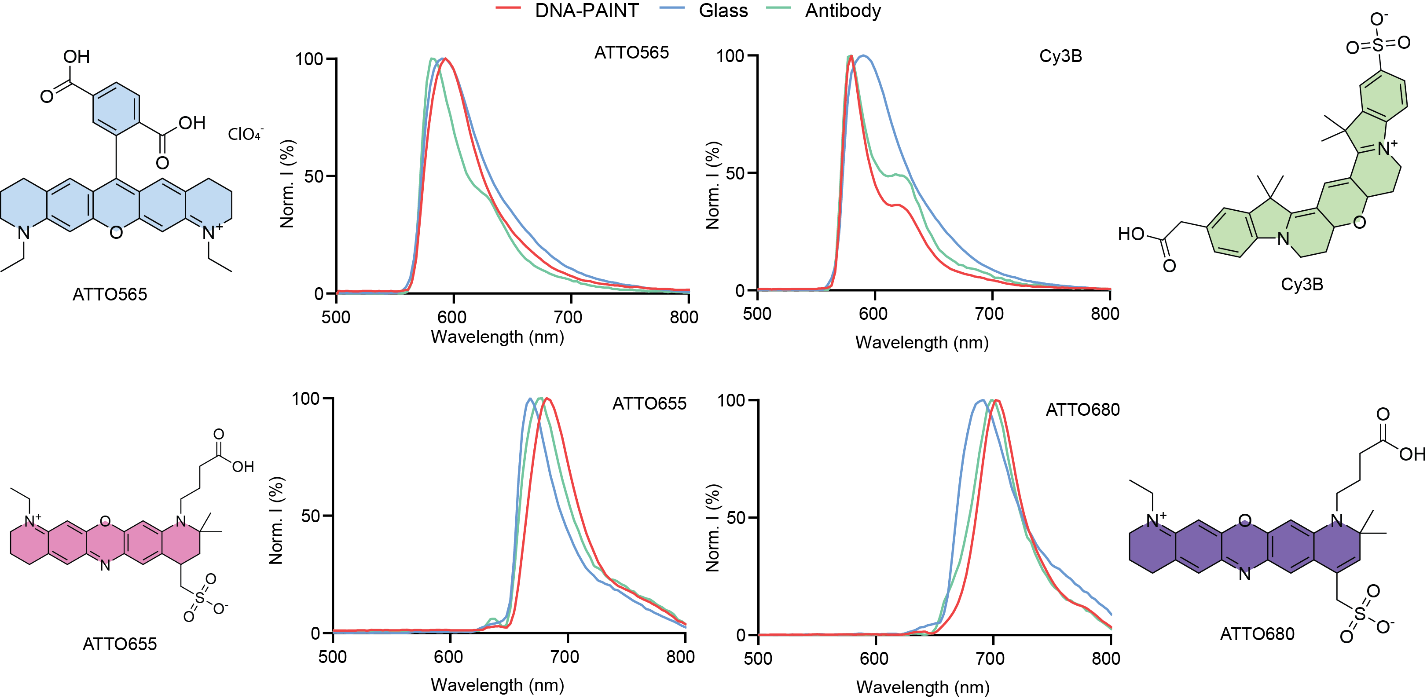

**Figure S4.** Averaged single-molecule emission spectra and molecular structures of ATTO565, Cy3B, ATTO655 and ATTO680 measured under DNA-PAINT (red curves), Glass (blue curves) and Antibody-conjugation (green curves) conditions and the respective ensembled emission spectra measured in aqueous solution at ~1-uM concentrations (black curves).

### 4. Imaging Acquisition and Analysis Procedure for multiplexed sDNA-PAINT

#### 4.1 Nanoruler

The nanoruler samples (GATTA-PAINT 80R, GATTAQUANT) were ordered from Massive Photonics and prepared according to manufacturer’s procedure and mounted on our sSMLM system and acquired either in the smFLUSH mode with a narrow entrance slit or without the imaging spectrometer to investigate the imaging condition. The kit includes all probes and reagents for washing and imaging.

##### 4.2 Multiplexed sDNA-PAINT imaging of BSC-1 cells

The fixed BSC-1 cell samples were customized ordered from Massive Photonics based on Massive-PAINTG 2-PLEX kit with its mitochondria and vimentin labeled with two orthogonal docking strands (F1 docking strand for vimentin and F2 docking strand for mitochondria) and imager strands. kit includes all probes and reagents for washing and imaging. Imager strands for F1 were labeled with ATTO680 and Cy3B; Imager strands for F2 were labeled with ATTO565 and ATTO655.

##### 4.3. Imaging acquisition and processing

The spectroscopic response of this imaging spectrometer was calibrated with a fluorescence lamp before each imaging session; the spectral dispersion is ~4 nm/pixel. The single-molecule images were acquired with a camera exposure time of 150 ms and 20,000 frames per acquisition. After the acquisition, the single-molecule images were processed with customized codes in MATLAB 2023b. For the sDNA-PAINT image, each single molecule is color-coded with its intensity-weighted spectra centroid.
